# Synergistic cytotoxicity with Chk1/Chk2-inhibitor prexasertib in small cell lung cancer following lurbinectedin-induced G2/M-checkpoint activation

**DOI:** 10.64898/2026.08.02.742299

**Authors:** Ashley F. Sanchez Sevilla Uruchurtu, Audrey Y. Su, Harshita Ganga, Shengliang Zhang, Ameen Raissi, Kevin Kwon, Tej Tummala, Tyler J. Roady, Jacqueline Moreno, Patrycja Dubielecka-Szczerba, Christopher G. Azzoli, Wafik S. El-Deiry

**Affiliations:** Pathobiology Graduate Group, Brown University, Providence, RI, USA; Laboratory of Translational Oncology and Experimental Cancer Therapeutics, Department of Pathology and Laboratory Medicine, The Warren Alpert Medical School of Brown University, Providence, RI, USA; Legorreta Cancer Center at Brown University, Providence, RI, USA; Department of Biomedical Engineering, Pennsylvania State University, University Park, PA, USA; Department of Biology, Johns Hopkins University, Baltimore, MD, USA; Zanvyl Krieger School of Arts and Sciences, Johns Hopkins University, Baltimore, MD, USA; College of Medicine, University of Central Florida, Orlando, FL, USA; Burnett School of Biomedical Sciences, University of Central Florida, Orlando, FL, USA; Hematology-Oncology Division, Department of Medicine, Brown University Health and The Warren Alpert Medical School of Brown University, Providence, RI, USA

**Keywords:** SCLC, lurbinectedin, cancer therapy, resistance mechanism, Chk1, Chk2, cell cycle checkpoint, prexasertib

## Abstract

Small cell lung cancer (SCLC) is an aggressive thoracic malignancy with a 5-year survival rate under 7%. Lack of meaningful improvement of survival rates despite advances in treatment highlights the need for novel therapeutic approaches to improve patient outcomes. Currently, carboplatin + etoposide chemotherapy is the backbone of treatment for most patients. Lurbinectedin is a cytotoxic drug with unique activity against small cell lung cancers in patients with extensive disease and acquired resistance to carboplatin + etoposide. Our preliminary experiments in human SCLC cell lines treated with lurbinectedin demonstrated a dose-dependent increase in Chk1 and Chk2 protein phosphorylation. A consequence of the frequent *TP53* inactivation in SCLC is tumor cell reliance on G2/M cell cycle checkpoints involving Chk1/Chk2 to maintain genomic integrity and allow cell survival following DNA damage. We hypothesised that inhibition of Chk1/Chk2-dependent responses with dual-inhibitor prexasertib (ACR-368), would potentiate tumor cell killing by lurbinectedin potentially in a synergistic manner. SCLC cells underwent cell death following single agent prexasertib exposure and this further increased with prexasertib + lurbinectedin combination. Highest Single Agent (HSA) synergy score calculations based on cell viability measurements suggested synergistic action between prexasertib and lurbinectedin at select dose combinations. Western blot analysis of intracellular proteins from SCLC cells treated with both drugs demonstrate dynamic, dose-dependent effects on Chk2, Chk1 and downstream effector Wee1, with lurbinectedin increasing intracellular levels of pChk1 and pChk2, while co-treatment with prexasertib deregulates this process across multiple human-derived cell lines. Synergistic killing was associated with elevated γ-H2AX levels indicative of DNA double strand breaks and PARP-cleavage due to apoptotic caspase activation. Despite some heterogeneity among treated SCLC cells, the increased phosphorylation of Chk1 was noted at several kinase-activating sites including Serine 296, 317, and 345 while Chk2 Tyrosine 68 phosphorylation was consistently upregulated by lurbinectedin. The results provide a preclinical mechanistic rationale for overcoming a pro-survival, drug resistance-promoting checkpoint pathway to enhance the unique efficacy of single-agent lurbinectedin in patients with SCLC.

## Introduction

Lung cancer is the leading cause of cancer death in both men and women. Small cell lung cancer (SCLC) is an aggressive neuroendocrine malignancy strongly associated with tobacco smoking. SCLC accounts for approximately 15% of new lung cancer cases and has a 5-year survival rate that remains under 10% despite recent advances in treatment^1,2^. First-line therapy for SCLC has traditionally comprised carboplatin + etoposide dual therapy with high response rates; however, relapse rates are high and are often accompanied by treatment resistance. Attempts at dose intensification, up to and including autologous stem cell transplantation to overcome dose-limiting neutropenia, failed to overcome mechanisms of resistance, or improve patient outcomes^3^. Adding immune-checkpoint therapies to standard chemotherapy results in limited improvement, with 12-months median overall survival versus 10 months with chemotherapy only for combination with atezolizumab or durvalumab^4,5^. In terms of novel immune therapies, the DLL3-targeted T-cell engager, tarlatamab, is more effective than chemotherapy as second-line treatment, and there are several antibody-drug conjugates in Phase II and III development, some with the potential for commercial approval based on Phase II data^6,7^.

Even with the rise of novel immune therapies, lurbinectedin remains a drug of both clinical and biologic interest because of its unique activity in small cell lung cancers which have become resistant to carboplatin + etoposide. In a Phase II study, lurbinectedin demonstrated a response rate of 22% (11-37%) in patients with platinum-refractory cancers, which likely accounts for its efficacy when integrated into first-line chemotherapy (started immediately after completion of carboplatin + etoposide)^8,9^. However, even with the addition of lurbinectedin to first line therapy, median survival is only 13 months. There therefore remains an unmet need for therapeutic approaches that meaningfully improve patient prognosis. SCLC’s propensity for treatment-resistant relapse and metastatic spread presents a particularly significant barrier to improving overall survival, necessitating the development of therapies designed to overcome mechanisms of resistance.

The DNA damage response (DDR) is one potential mechanism whereby tumor cells evade complete eradication by chemotherapy. In healthy cells, checkpoint mechanisms monitor damaged or improperly duplicated DNA to maintain genomic integrity and repair damaged DNA prior to cell division. However, these same mechanisms are used by cancer cells to evade mitotic catastrophe following administration of DNA-damaging agents^10^. Recent pre-clinical studies and clinical trials have explored possible druggable targets in the DDR machinery, including ATR, ATM and checkpoint kinases (Chk) 1 and 2^11–15^. The success of these trials has been limited, with off-target cardiac toxicity as a concern along with significant bone marrow and gastrointestinal toxicities.

In terms of mechanism of action, lurbinectedin is a small molecule DNA damaging agent that exerts its genotoxic effects by targeting the DNA strand in two ways. Lurbinectedin binds RNA Polymerase II, causing DNA replication-fork collapse (RFC) and it also binds directly to the minor groove of the DNA double helix to produce double-stranded breaks (DSB)^16,17^. The first mechanism of action activates the Ataxia Telangiectasia and Rad3-related protein (ATR)-mediated arm of the DDR, activating downstream effector Chk1 and its substrates Wee1 and cdc25A. Meanwhile, DSBs recruit and activate Ataxia-telangiectasia mutated protein (ATM), which subsequently phosphorylates and activates Chk2 (**Figure 1**) This dual response to a single drug makes lurbinectedin the ideal candidate to pair with a checkpoint inhibitor, to evaluate the combined effects and parse out redundancies between each arm of the DDR.

**Figure 1.**
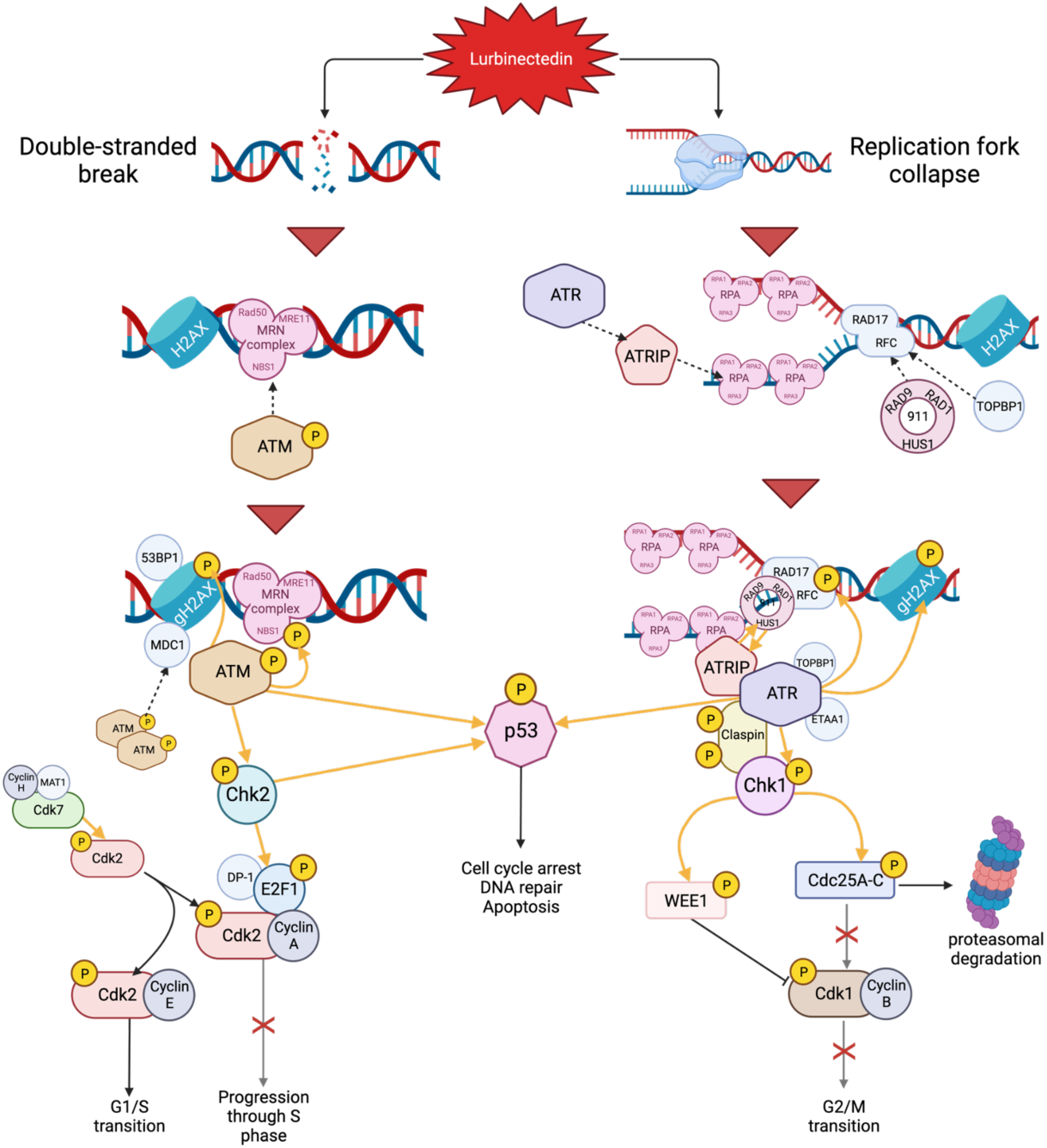
DSBs and RFC by lurbinectedin activate ATM/ATR mediated DNA damage repair. Lurbinectedin induces DNA damage by acting as an alkylating agent, binding the minor groove of the DNA double helix, forming double-stranded breaks (DSBs). Lurbinectedin also irreversibly stalls RNA Polymerase II, which causes replication fork collapse (RFC). DSBs recruit and activate ATM, which in turn recruits Chk2 and converts it into its active state. RFC mediates activation of Chk1 by way of ATRIP and ATR. In TP53-competent cells, ATM and ATR also modulate p53 activation, inducing cell cycle arrest, DNA repair, and/or apoptosis. Tumor cells where p53 is inactivated, such as in small cell lung cancer, then rely on Chk1/2 to prevent entry into mitosis in the event of DNA damage.

Chk1 is a key regulator of cell cycle and cell survival. Chk1 activation (**Figure 1**) bridges the DNA damage response and cell cycle checkpoints. Activation of Chk1 results in the initiation of the checkpoint, triggering cell cycle arrest, RNA repair and, if repair is not possible, controlled cell death^18^. Active Chk1 impacts various stages of cell cycle: S phase^19^, G2/M transition^20^, and mitosis^21^. In S phase, Chk1 is essential for maintenance of genomic integrity. It monitors DNA replication and responds to genotoxic stress, stalling DNA replication to give time for DNA repair mechanisms to restore the genome. During the G2/M transition, Chk1 acts as a signal transducer between DNA damage and G2/M checkpoint activation. Activation of the checkpoint arrests the cell in G2 phase until its DNA is repaired, and the cell is ready for mitosis. Apoptosis occurs if the damage is irreversible. Chk1 inhibits G2/M transition by preventing formation of the cdk1/cyclin B complex. Phosphorylation of nuclear kinase Wee1 by pChk1 inhibits Cdk1 activation. Chk1 also phosphorylates cdc25C, leading to its proteosomal degradation, thereby preventing dephosphorylation of cyclin B-bound cdk1. Chk1 must inactivate for the cell to enter M phase.

Chk1 has been found to be overexpressed in many tumor types, including breast, colon, and liver cancer. Levels of Chk1 expression correlate with tumor grade and disease recurrence, which suggests it may have a role in promoting tumor growth by conferring resistance to DNA damaging agents. Studies have demonstrated that inhibiting Chk1 reactivates tumor suppressive activity of the protein phosphastase 2A (PP2A) complex^22^. Chk1 may have a particularly crucial role in SCLC survival. Tumors with *TP53* deficiencies, such as SCLC, rely heavily on Chk1-mediated cell cycle arrest to maintain genomic integrity^23^. This dependence on checkpoint machinery makes SCLC a promising target for DDR effectors blockade.

Chk2 acts as a signal transducer in response to DSBs and has dual roles in cancer initiation and progression (**Figure1**). Chk2 is activated through phosphorylation by ATM, a sensor of DNA damage^24^. Active Chk2 phosphorylates cell cycle transcription factors E2F1 and PML. Chk2 is involved in the regulation of DNA repair, cell cycle arrest and/or apoptosis in response to DNA damage^25^. Its role in tumor suppression arises from its ability to mitigate rapid and uncontrollable cell division in pre-cancerous cells. In TP53-competent cells, Chk2 can also stabilise the p53 protein, leading to cell cycle arrest in phase G1^26^. Despite CHEK2’s role as a suppressor of tumor initiation, Chk2 plays a very different part in cancer cell survival in CHEK2 wild-type tumors. In cancer patients with functional Chk2 protein, activated Chk2 acts in a similar manner to Chk1, mediating the DDR and allowing the tumor cells to evade mitotic catastrophe^27^. Potential synthetic lethality approaches correlate with *CHEK2* state; in patients with *CHEK2* mutated tumors, inactivation of other DDR proteins such as poly(ADP-ribose) polymerase (PARP) in conjunction with CHEK2 loss of function may enhance cancer cell killing. Conversely, inhibiting activity of wild-type Chk2 protein in tumors with mutations in other DDR genes, or in combination with DNA damaging therapies, could similarly potentiate tumor cell death^28^

Prexasertib (ACR-368).is a small molecule checkpoint kinase inhibitor that is primarily active against Chk1 and has minor activity against Chk2. Prexasertib preferentially binds Chk1, preventing autophosphorylation at S296 and catalytic activation. There is evidence that prexasertib may disrupt AMP-activated protein kinase (AMPK) kinase activity^29^. In a phase 2 study, prexasertib showed little or no single-agent activity in patients with previously-treated small cell lung cancer, and caused high rates of neutropenia^30^. When considering neutropenia as a side effect of therapy, it is important to note that Trilaciclib, an intravenous CDK 4/6 inhibitor, is in routine clinical use in patients with ES-SCLC, specifically because it prevents cytopenias following treatment with carboplatin + etoposide by providing a transient, protective G1 cell cycle arrest in proliferating bone marrow cells^31^.

In the present study, we aimed to address the need for a therapeutic approach that both targets the DDR and potentially reduces off-target effects. To this end, we combined lurbinectedin, an FDA-approved treatment for ES-SCLC, with Chk1/2 dual inhibitor prexasertib. These preclinical studies document activation of the Chk1-dependent G2/M checkpoint by lurbinectedin, and demonstrate the potential of incorporating Chk1/Chk2 inhibitors such as prexasertib to overcome this cell survival, drug-resistance mechanism.

## Results

### Treatment of SCLC cell lines with lurbinectedin induces activation of DDR effector proteins and markers of apoptosis in a dose-dependent manner

We explored the molecular consequences of lurbinectedin treatment of human SCLC cells to better understand its mechanism of action as well as potential drug resistance mechanisms. Histone protein H2AX can act as an early sensor of DNA damage. At sites of DSB, changes to chromatin structure promote phosphorylation of H2AX by ATM and ATR. γH2AX, the phosphorylated form of H2AX, then recruits additional proteins involved in initiation of the DNA damage checkpoint (**Figure 1**)^32^. Constitutive levels of γH2AX in untreated cells are believed to represent DNA damage by endogenous oxidants^33^. The dose-dependent increase of γH2AX observed in treated SCLC cells compared to controls indicates accumulation of DNA damage in response to lurbinectedin (**Figure 2**).

**Figure 2.**
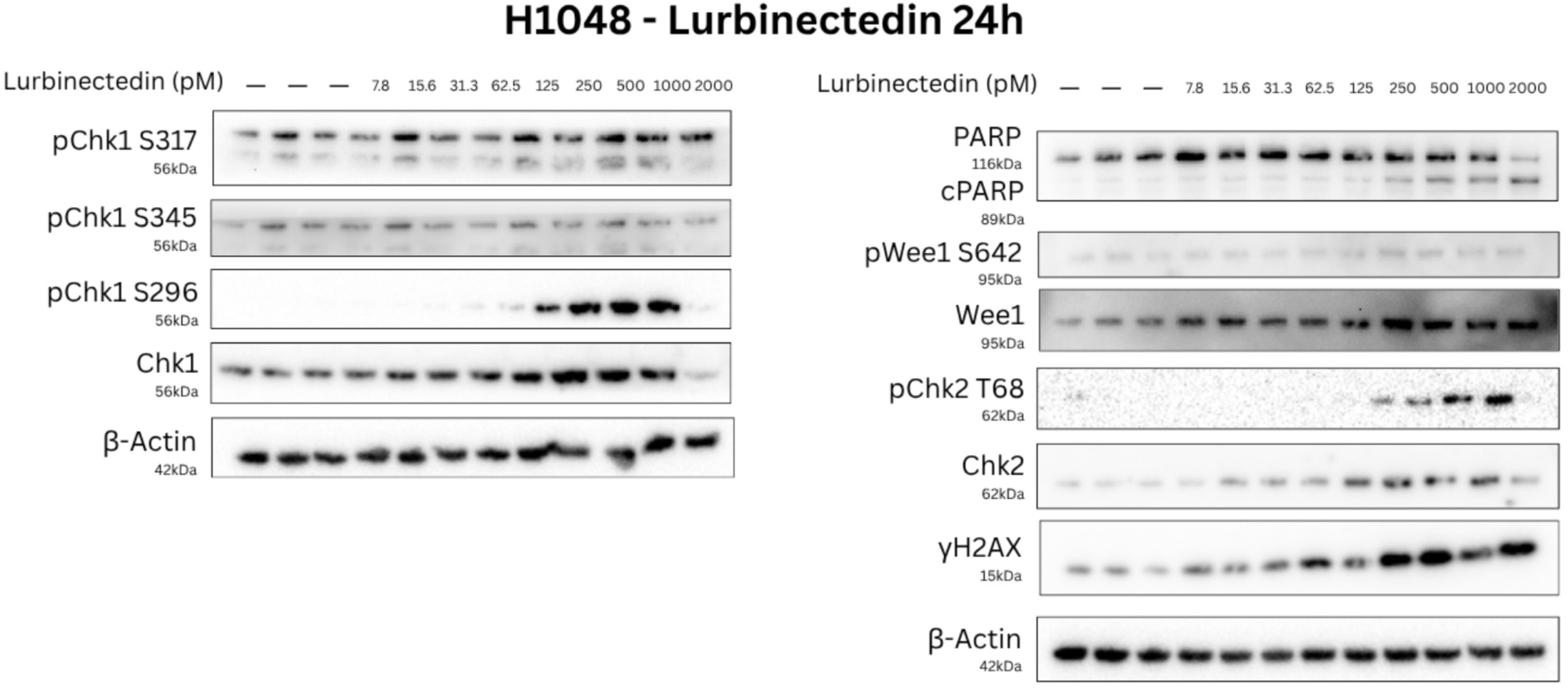
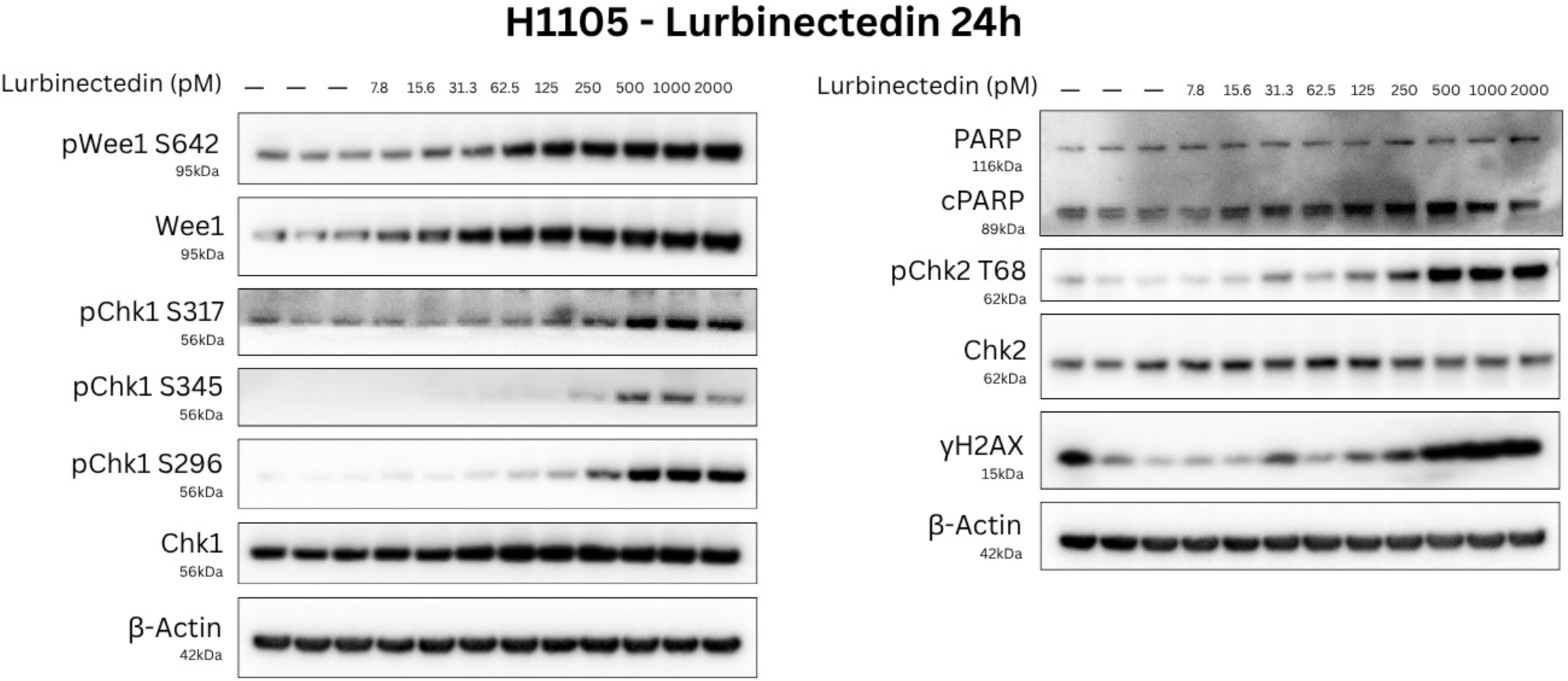
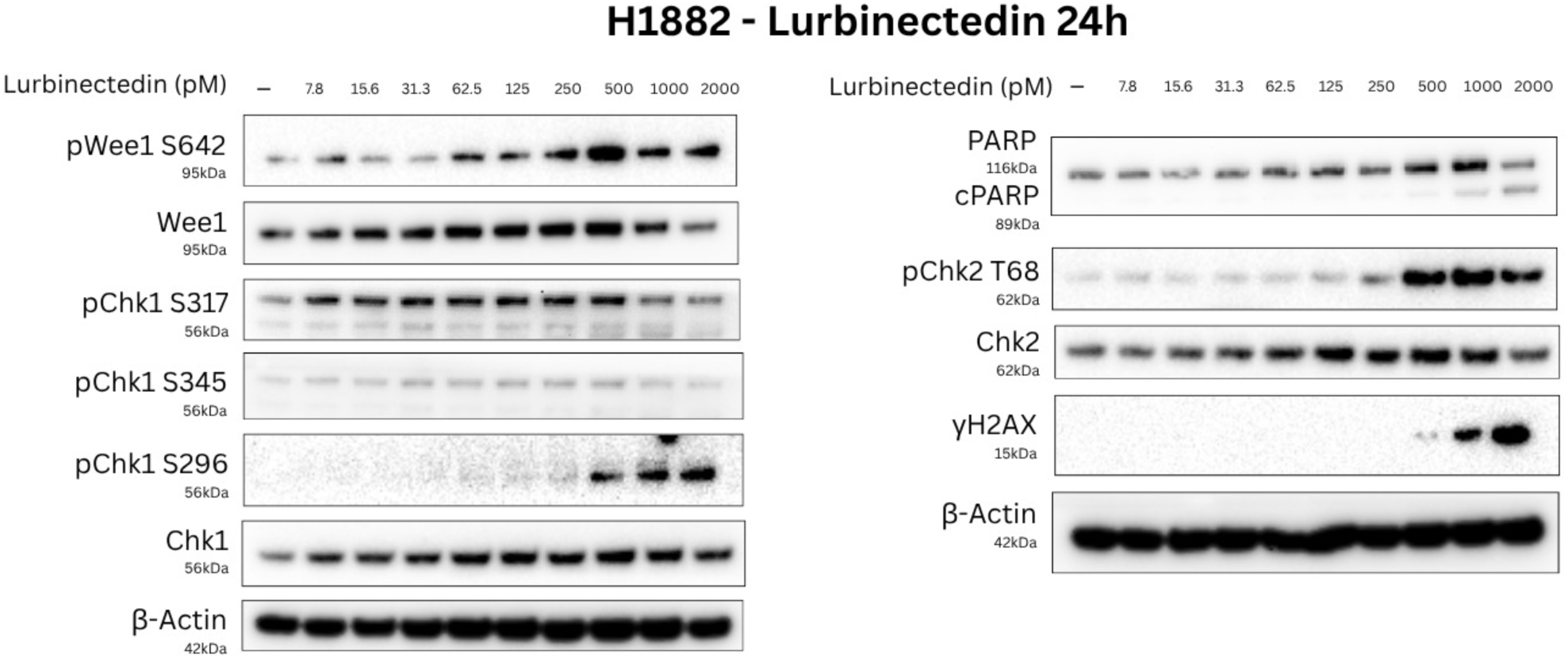
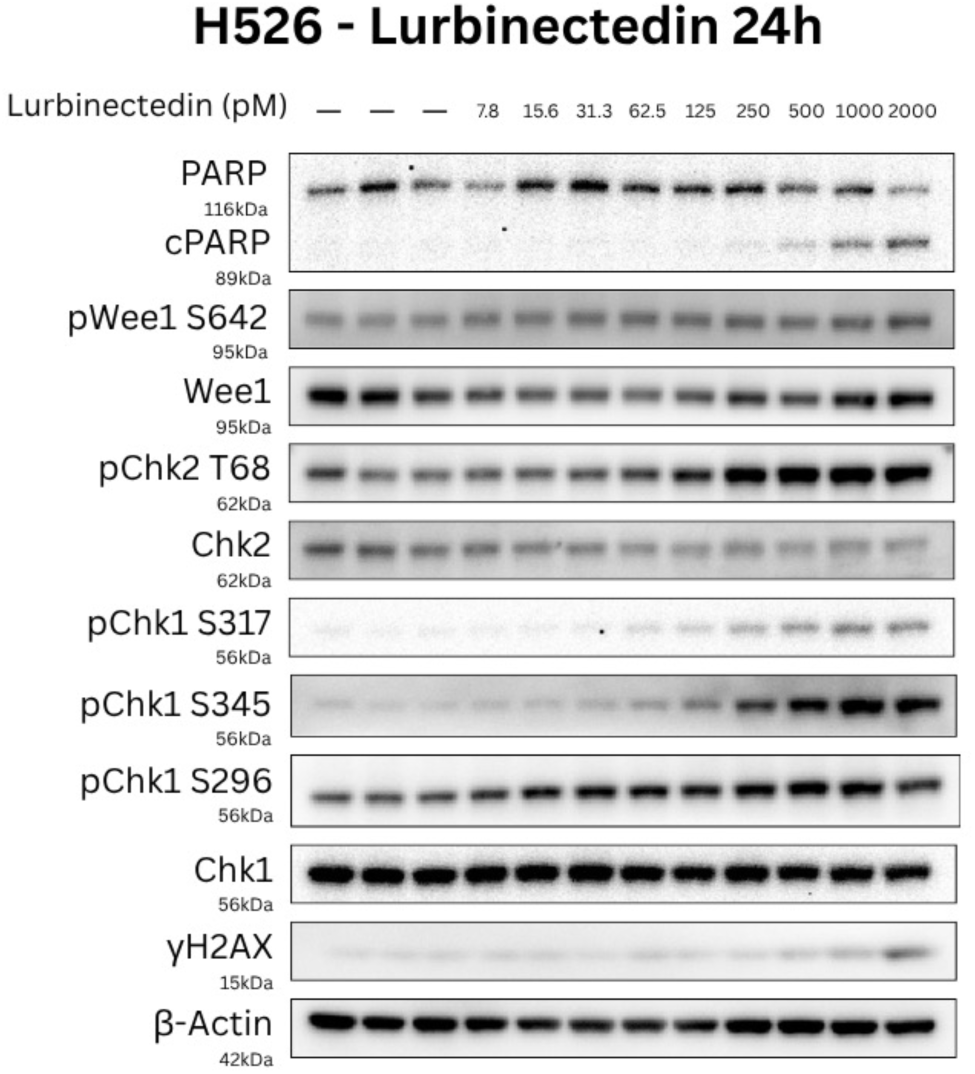
Treatment of SCLC cell lines with lurbinectedin induces activation of DDR effector proteins and markers of apoptosis in a dose-dependent manner. A-D Four human-derived SCLC cell lines (NCI-H1048 (ATCC CRL-5853), NCI-H1105 (ATCC CRL-5856), NCI-H1882 (ATCC CRL-5903), NCI-H526 (ATCC CRL-5811)) were treated with lurbinectedin for 24h. The cells were then harvested and their protein content extracted for analysis by immunoblotting.

Nuclear PARP1 is an enzyme involved in various cellular processes. Notably, it is involved in regulating tumor cell proliferation and differentiation, as well as DNA repair. PARP1 is an early sensor of DNA damage and coordinates the initiation of base excision repair in response to ssDNA breaks. PARP1 has additional functions in the regulation of cell death. Depletion of cellular ATP by activated PARP can lead to necrotic cell death^34^. PARP1 can also trigger apoptosis in a caspase-dependent manner by stimulating the mitochondria to release Apoptosis Inducing Factor (AIF)^35^. Cleavage of PARP by caspases subsequently inactivates PARP. Therefore, PARP cleavage forming an 89kDa fragment can be measured by immunoblotting and used as a measure of apoptosis. In all four cell lines we noted an increase in this 89kDa cPARP fraction following administration of lurbinectedin, indicating induction of programmed cell death (**Figure 2**).

### Lurbinectedin alters phosphorylation states of Chk1 and Chk2 in a dose dependent manner

In results from Western blot experiments, we saw dose-dependent effects on Chk1 phosphorylation in all four cell lines tested; however, results for each cell line differ (**Figure 2**). In H526 cells, treatment with lurbinectedin robustly increased pChk1-S345 in a dose-dependent manner (**Figure 2D**). H1105 cells exhibited an increase in pChk1-S345 correlating with increasing drug doses, followed by a decrease at the highest dose (**Figure 2B**). By contrast, H1048 and H1882 cells demonstrated low-level constitutive levels of pChk1-S345, which were slightly reduced by lurbinectedin at the highest doses (1-2 nM) (**Figures 2A, 2C**).

As with pChk1-S345, immunoblotting of H1048 cells revealed the presence of pChk1-S317 in the control samples, which was overall unaffected by administration of lurbinectedin (**Figure 2A**). pChk2-S317 levels in H1105 rose in response to lurbinectedin but did not decrease at the highest dose, unlike pChk2-S345 (**Figure 2B**). H1882 cells also constitutively expressed pChk1-S317; this phosphorylated form of Chk1 declined at 1nM and 2nM concentrations of lurbinectedin (**Figure 2C**). Finally, pChk1-S317 in H526 responded to lurbinectedin concurrently with pChk1-S345 (**Figure 2D**).

pChk1-S296 expression in H1048, H1104, an H1882 was elevated by lurbinectedin in a dose-dependent manner (**Figure 2A-C**). Wee1 phosphorylation, on the other hand, was not equal between cell lines despite the consistent activation of pChk1-S296: in H1048, levels of pWee1 and total Wee1 did not appear to be affected by lurbinectedin. H1105 cells demonstrated dose-dependent increases in quantities of pWee1 and total Wee1. H1882 cells appear to upregulate Wee1 phosphorylation in response to lurbinectedin while levels of total Wee1 counterintuitively wane. By comparison, lurbinectedin treatment of H526 did not significantly increase pChk1-S296 or pWee1 (**Figure 2D**). Chk2 phosphorylation at T68 was consistently upregulated by lurbinectedin across all cell lines (**Figure 2**).

### Prexasertib synergises with lurbinectedin in SCLC cells treated at certain dose combinations

Cell viability data at 72 hr following combination treatment was used to calculate synergy between lurbinectedin and prexasertib. To evaluate synergy, the drug-response matrix was compared with scoring models assuming no interaction (null-models). Deviation of observed data from the response anticipated by the null-model was used to calculate synergy, additive effect, or antagonism. Based on the known mechanisms of actions for lurbinectedin and prexasertib (i.e. two drugs independently enacting their effects within the same response cascade in a non-stochastic pathway), we opted to use the highest single agent (HSA) reference model, which equates the expected combination effect with the maximum responses to each drug at the given concentrations. HAS is most applicable when seeking to characterise potentiation– i.e., a condition in which a given drug increases the effect of another drug when administered concurrently^36^. Thus, the synergy score calculated using this model (S_HSA_) can be expressed as: S_HSA_ = E_A,B,…,N_ - max(E_A_, E_B_,…, E_N_), where EA,B,…,N is the combined effect of N drugs and EA, EB,…EN) are the responses of each single drug. Using this model, we calculated synergy scores for lurbinectedin-prexasertib double treatment in four cell lines– H1048, H1105, H1417, and H1882 (**Figure 3A-C**). Overall findings indicate low-level synergy (HAS scores <10), with higher individual synergy scores at specific dose combinations (H1048: 62.5 pM lurbinectedin + 3.125 nM prexasertib, H1105: 125 pM lurbinectedin + 62.4 nM prexasertib, H1882: 31.3 pM lurbinectedin plus 12.5 nM prexasertib, H1417: 250 pM lurbinectedin plus 100 nM prexasertib).

**Figure 3.**
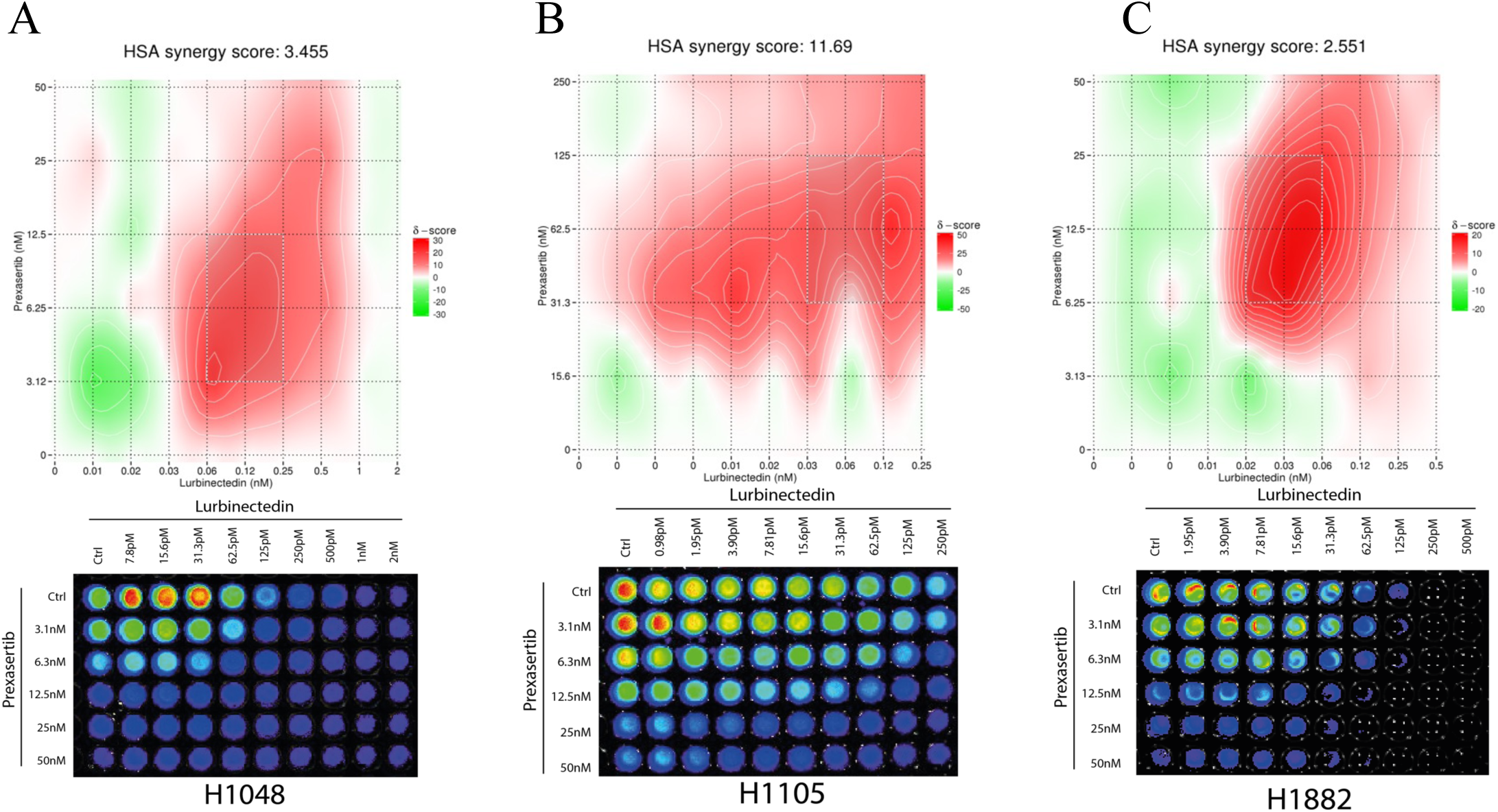
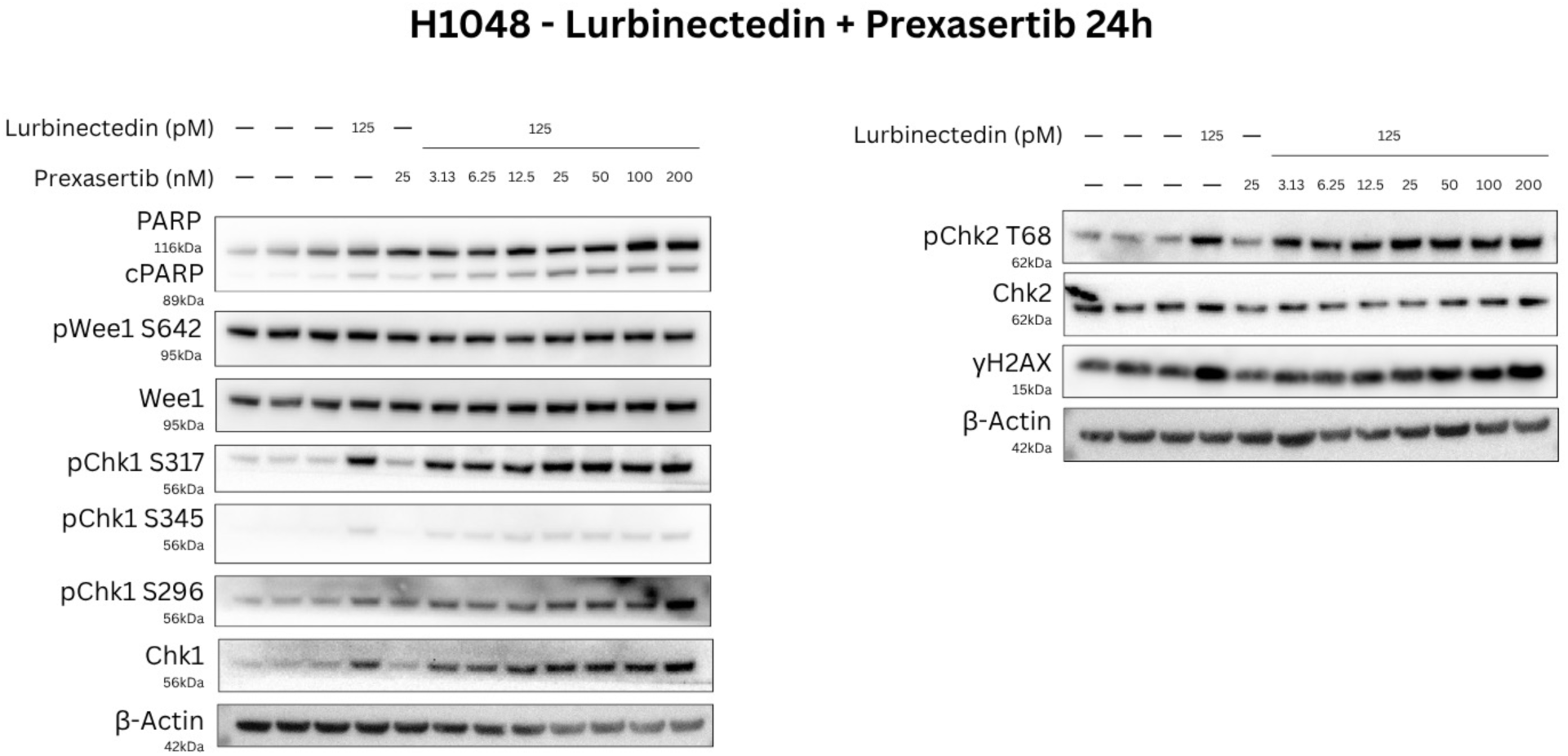
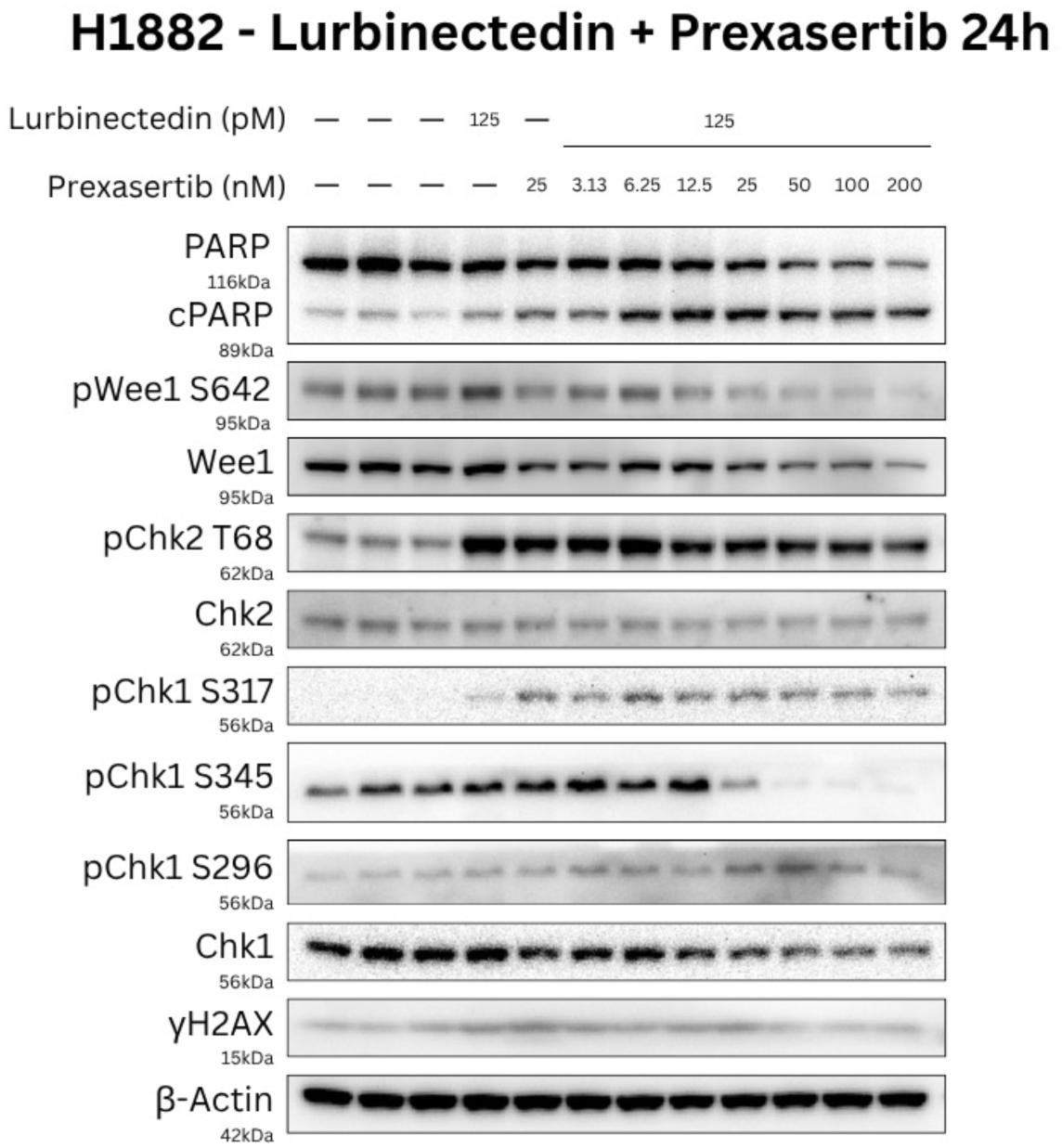
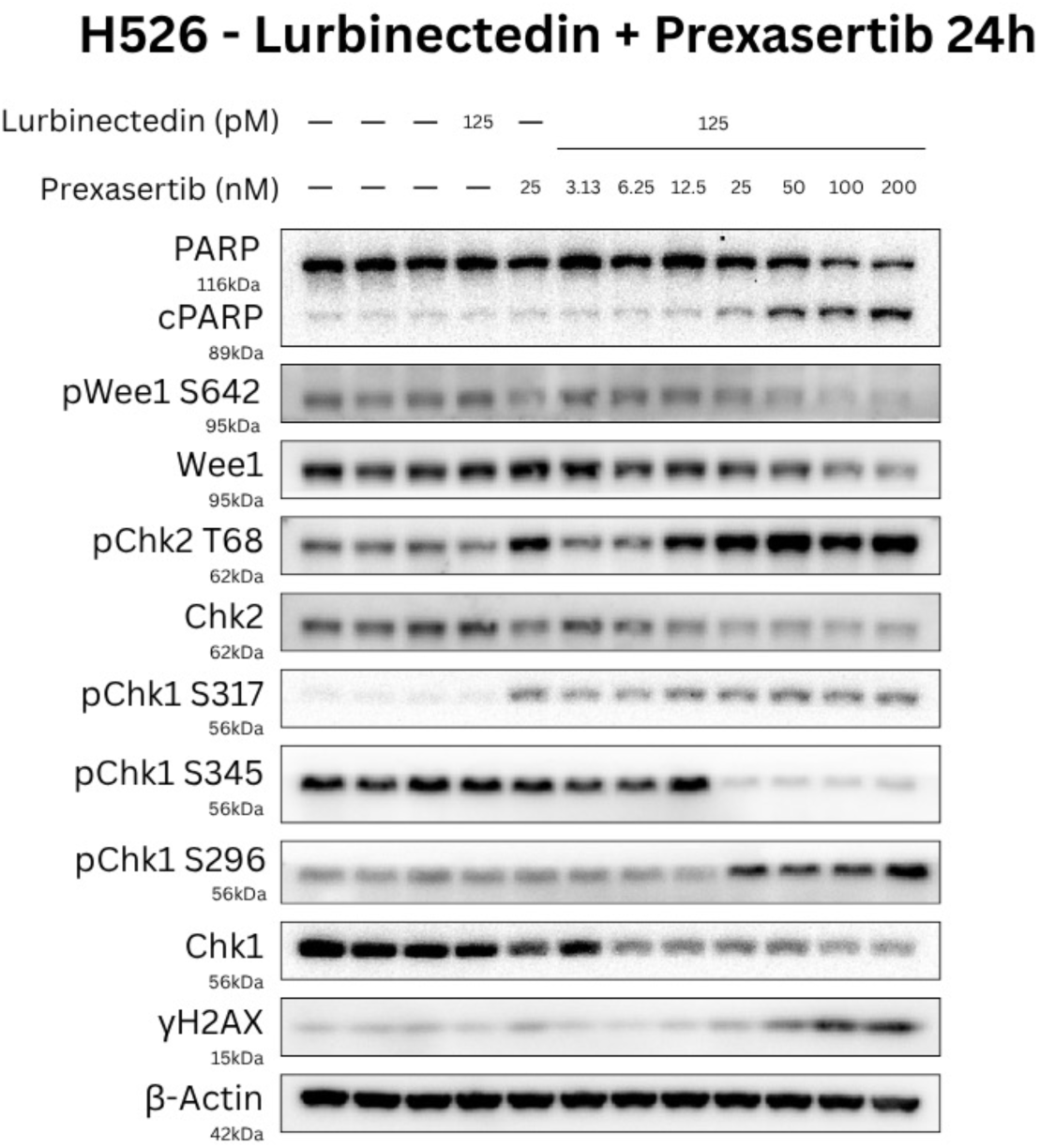
Prexasertib potentiates the genotoxic and cytotoxic effects of lurbinectedin and dysregulates Chk1/Chk2 activation. A-C: Three of our SCLC cell lines (NCI-H1048, NCI-H1105, NCI-H1882) were treated with lurbinectedin and prexasertib at serial doses. Cell viability was assessed at 72h by CellTiter Glo; the quantitative luminescence reading output data was entered into SynergyFinder to calculate drug synergy by comparing cell death relative to controls. D-G: SCLC cell lines were treated with lurbinectedin, prexasertib, or combination 24h. The cells were then harvested and their protein content extracted for analysis by immunoblotting.

Prexasertib potentiates the genotoxic and cytotoxic effects of lurbinectedin and dysregulates Chk1/Chk2 activation

When H1048 was treated with prexasertib alone, there was no visible difference in Chk1 phosphorylation at S345 compared with controls. However, when combined with lurbinectedin, we observed an increase in pChk1-S345 (**Figure 3D**). In H1882 and H526 we noted an inverse effect (**Figure 3E, F**): prexasertib abrogated the phosphorylation of S345 by lurbinectedin shown in **Figure 2C and 2D**. Concentrations of pChk1-S317 in all cell lines were higher in combination treated samples than in the controls. Increasing the dose of prexasertib co-administered with lurbinectedin had no appreciable effect (**Figure 3D-F**). Chk1 phosphorylation at S296 was increased by prexasertib in combination with lurbinectedin. As with lurbinectedin monotreatment, levels of total and phosphorylated Wee1 were unaffected in H1048 when prexasertib was added (**Figure 3D**). The upregulation of pWee1 and total Wee1 in H1882 induced by lurbinectedin was abrogated in a dose-dependent mode by prexasertib (**Figure 3E**). Despite H526 cells’ lack of response to lurbinectedin in terms of absolute Wee1 and its phosphorylation, addition of prexasertib did reduce constitutive levels of both forms of Wee1 (**Figure 3F**). In H1048 cells, prexasertib did not reduce pChk2-T68 levels when combined with lurbinectedin at the dose range used, whereas prexasertib abrogated Chk2 phosphorylation in a dose-dependent manner in H1882 (Fig. 3D, E). In H526, the combination of prexasertib with lurbinectedin increased pChk2-T68 in a dose-dependent manner while paradoxically reducing total Chk2 availability (**Figure 3F**). When H526 and H1048 cells were treated with lurbinectedin and prexasertib together, γH2AX was elevated at the highest doses of prexasertib, indicating increased levels of DNA damage accumulation versus lurbinectedin alone (**Figure 3D, F**). In H1882, we did not note an elevation of γH2AX in response to the treatment doses used (**Figure 3F**). None of our cell lines demonstrated significant activation of γH2AX in response to prexasertib alone. When each of the SCLC cell lines were treated concomitantly treated with prexasertib and lurbinectedin, we noted a significant increase in PARP cleavage as compared to lurbinectedin-only treated samples (**Figure 3D-F**).

Combination lurbinectedin plus prexasertib treatment of synchronized H1048 SCLC cells for 24 hr is associated with a higher proportion of cells with sub-G1 DNA content versus control or monotherapy

DNA fragmentation by endonuclease activity is a feature of late-stage apoptosis. The resulting fragments accumulate in cells. By permeabilising the cells and washing in phosphate-citrate buffer, these fragments leak out, leaving behind a population of cells with decreased DNA content. A quantitative DNA profile of a given sample of cells can therefore be calculated using a nucleic acid stain like propidium iodide (PI) by way of flow cytometry. The resulting histogram can be interpreted to calculate the populations of cells at each stage of the cell cycle within the sample^37^. Apoptotic cells with depleted DNA are represented in the region left of the G1 peak, i.e. sub-G1 (**Figure 4**). To assess effects of single- and double-agent therapy on apoptosis and cell cycle accumulation, we synchronised cells by serum deprivation followed by serum release. Cells were treated with lurbinectedin, prexasertib, or combination for 24h immediately on release. Each sample was then fixed in EtOH and stained with PI for analysis by flow cytometry. We observed a higher proportion of cells in the subG1 region of the histogram when cells were treated with lurbinectedin and prexasertib together, in comparison to the control group and the single-agent treatment groups (**Figure 4**). The results of our analysis are in concordance with previous published studies including Zhang *et al.*^38^ who noted similar patterns of cell cycle distribution in their 2026 study that combined cisplatin with prexasertib in hypopharyngeal squamous cell carcinoma cell lines.

**Figure 4.**
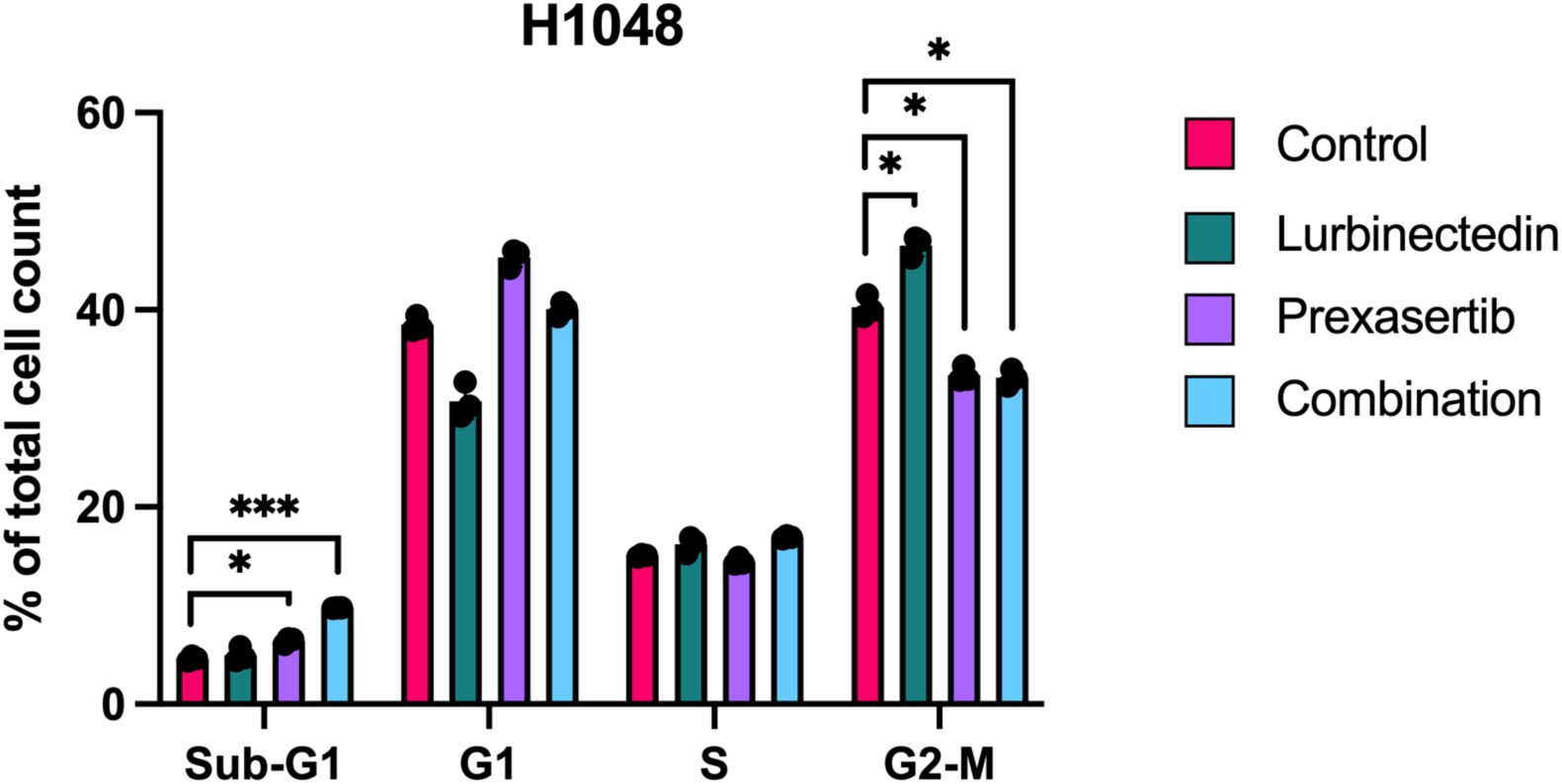
treatment with lurbinectedin and prexasertib exert cytotoxic effects at different stages of the cell cycle. H1048 cells were synchronised by serum deprivation for 48 (0.5% FBS HITES). Cells were then released in standard 5% FBS HITES media and treated with lurbinectedin, prexasertib, or combination for 24h. Cells were then fixed in 70% EtOH and stained in propidium iodide for Sub-G1 analysis by flow cytometry.

Aurora B, a serine-threonine kinase, mediates chromosome segregation by regulating the orientation of sister chromatids during metaphase and anaphase. Overexpression of AURKB induces aneuploidy and is often found in human carcinomas. Inhibition of Aurora kinase B by small molecule inhibitor BI 811283 was shown to override the metaphase to anaphase transition checkpoint, resulting in genomic instability, senescence, and increased cell death^39,40^. In order to evaluate the effect of lurbinectedin and prexasertib on Aurora B expression, human SCLC cell lines were treated with lurbinectedin, prexasertib, or both in combination (**Supplementary Figure 2**). Lurbinectedin increased availability of Aurora B protein in a dose-dependent manner, while treatment with prexasertib abrogated endogenous levels of the kinase. When treated with concurrently increasing doses of both drugs, Aurora B levels remained stable.

Effects of treatment on availability of Aurora kinase B may shed further light on how lurbinectedin and prexasertib affect cell cycle progression in SCLC. Sub-G1 analysis of cell line H1048 indicated that a higher proportion of cells enter G2/M when treated with lurbinectedin versus control. This is supported by our observation that lurbinectedin increases levels of Aurora B (**Supplementary Figure 2**), which is responsible for mediating the metaphase/anaphase transition checkpoint. Conversely, prexasertib reduced levels of Aurora B, which, in conjunction with the higher proportion of Sub-G1 cells observed in **Figure 4**, suggests disruption of cell cycle checkpoints and increased cell death.

There are certain limitations to this method as gating for sub-G1 doesn’t account for all dead cells. Necroptosis secondary to genotoxic stress, for example by chemotherapy, can cause cells to accumulate in G2/M. This could explain the G2/M peak seen in the lurbinectedin-only treatment group. Therefore, there are limits to the utility of this method in calculating cell cycle stage in populations of cells exposed to different cytotoxic agents.

## Discussion

In this study, we have demonstrated synergy between the genotoxic agent lurbinectedin and small molecule checkpoint kinase inhibitor prexasertib in three human SCLC cell lines. We have further shown how these drugs affect activation of DDR effector proteins, both as single-agent treatments and in combination.

### SCLC cell lines respond differently to lurbinectedin and prexasertib

Immunoblotting results show that lurbinectedin, alone and in combination with prexasertib, affects Chk1 phosphorylation differently in each cell line despite equal dose and duration of treatment.

The S345 and S317 phosphorylation sites, located on the regulatory C-terminal domain of Chk1, are phosphorylated by activated ATR^41^. Phosphorylation at S345 is essential for DNA damage checkpoint activation (and therefore cell viability); it is also required for Chk1 catalytic activity and function, creating a docking site for 14-3-3β/ζ. 14-3-3β/ζ binding results in nuclear retention and accumulation^42^. This combination of activation and nuclear accumulation accelerates checkpoint activation in response to DNA damage. Following ATR-dependent phosphorylation of Chk1 at S345 and S317, Chk1 autophosphorylates at S296^43^, after which S345 and S317 are dephosphorylated by PP2A and Wip1^43,44^. pChk2 S296 can be detected across the entire nucleus, whereas pChk1 S345 is primarily detected at DNA damage foci; this suggests an epistatic model wherein S345 dephosphorylation and subsequent S296 autophosphorylation promotes release of Chk1 from the chromatin, allowing dissemination of signal. Cdc25A, a direct substrate of pChk1, is likewise distributed diffusely in the nucleus^45^; ergo, distribution of active Chk1 in the nucleoplasm is essential for its downstream functions.

In two of our cell lines, we observed dose-dependent lurbinectedin-induced increase in pChk1-S345 followed by decreased levels at 2 nM while pChk1-S296 increased uniformly. Given then that ATR-dependent phosphorylation of pChk1 at S345 is followed by autophosphorylation at S296 and dephosphorylation of S345, these results could indicate that as the dose of lurbinectedin increases, the phosphorylation shift from S345 to S296 may be happening at a faster rate. To assess this possibility, future experiments should collect intracellular proteins at the same doses of lurbinectedin but at shorter time-points. Results from these experiments together could then illustrate a time course of the Chk1 phosphorylation cascade in response to lurbinectedin at increasing doses.

A 2011 study on Chk1 inhibition in conjunction with gemcitabine in pancreatic cells could provide insight into the effects of prexasertib in conjunction with lurbinectedin on pChk1-S345^46^. The authors concluded that an observed increase in pChk1-S345 following administration of Chk1i AZD7762 was caused by accumulation of DNA damage, measured by relative levels of γH2AX. This observation supports our own findings in cell line H1048, where pChk1-S345 increases concurrently with γH2AX in the combination treatment group, and in H1882, where the opposite effects are seen. In H526, however, we see a reversal of this observation– while S345 phosphorylation decreases as prexasertib dose increases, γH2AX levels increase. In their paper, Venkatesha *et al*. noted that inhibition of PP2A, which dephosphorylates S345, also plays a role in pChk1-S345 increase. Functional analysis of PP2A protein in each cell line could therefore help elucidate the mechanism underpinning the differences seen.

When treating H526 cells with lurbinectedin, escalating the dose increased levels of pChk1-S317 and S345, but not pChk1-S296. In the same Western blot, we observed that accumulation of γH2AX was less pronounced than in the other three cell lines. This may suggest that reduced activation of H2AX, while sufficient to trigger ATR-mediated Chk1 phosphorylation, is perhaps insufficient to trigger phosphorylation of Chk1 at S296. This is supported by the observed lack of Wee1 phosphorylation, Wee1 being a direct substrate of activated Chk1. By comparison, in cell lines where pChk1-S296 increases, we also observed increases in pWee1. Another measure of pChk1-S296 activity is the phosphorylation and degradation of cdc25; by probing for cdc25C we could verify whether the observed lack of Wee1 phosphorylation is indeed caused by lack of pChk1 kinase function.

When H526 was treated with lurbinectedin and prexasertib together, γH2AX was elevated at the highest doses of prexasertib, indicating increased levels of DNA damage accumulation versus lurbinectedin alone (**Figure 3F**). We also observed increased levels of pChk1-S296, which supports the above conclusion that lack of Chk1-S296 phosphorylation in the lurbinectedin-only cells is secondary to insufficient DNA damage. Despite the increase in pChk1-S296 protein, its catalytic function appeared to be impaired by prexasertib, evidenced by decreased Wee1 phosphorylation and total Wee1 protein linked to increased drug dose.

Levels of pChk2-T68 were increased by lurbinectedin treatment in all four cell lines tested (**Figure 2A-D**). This is an expected result given lurbinectedin’s activity as an alkylating agent inducing DSB. When combined with prexasertib, results again diverged; in H1882, prexasertib reduced levels of pChk2-T68 (**Figure 3E**), in accordance with prexasertib’s anticipated function as a Chk2 inhibitor. In H1048, the administered doses of prexasertib seemed to be insufficient to overcome the activation of Chk2 by lurbinectedin (**Figure 3D**). In H526, prexasertib in fact seems to potentiate this activation (**Figure 3F**).

Further experiments are required to evaluate whether the changes to pChk2-T68 levels also affect its effector function; as described above, the most direct readout is to assess its substrates. If pChk2-T68 activity is diminished, we would expect to see a decrease in pE2F1; if its activity is increased, pE2F1 would also increase.

The question remains as to what intrinsic differences between these cell lines underpins the variability in response to the same genotoxic agents. Transcriptomic and proteomic analysis and functional assays of RNA Polymerase II and ATR would elucidate whether differences in the replication fork machinery upstream of Chk1 underlie the discrepancies observed. Similarly, transcriptional and functional evaluation of the proteins involved in ATM recruitment and activation could provide answers regarding the Chk2 response to prexasertib and lurbinectedin. More detailed readouts of Chk1/2 effector function and transcriptional profiling of the cell lines used here may help elucidate the precise mechanisms underpinning lurbinectedin-prexasertib synergy and the role of genetic variances in treatment efficacy.

### Prexasertib induces accumulation of DSBs in conjunction with lurbinectedin but not on its own

It has been demonstrated that prexasertib causes accumulation of DSBs as measured by γH2AX; however, it has not been definitively shown that prexasertib itself binds or directly damages the DNA. In the present results we observed that co-treatment with prexasertib increased γH2AX levels in a dose-dependent manner even when the dose of lurbinectedin remained the same. However, we did not observe evidence that prexasertib alone induced significant increases in γH2AX levels at the dose tested. Further testing with a wider range of prexasertib doses in SCLC cell lines may help assess its efficacy in disruption of DNA repair as a single agent.

### Alternative approaches to synthetic lethality with lurbinectedin

Knockdown of PARP1 by siRNA or with use of PARP inhibitors (PARPi) impairs repair of ssDNA breaks, sensitising cells to alkylating agents. When ssDNA breaks occur during DNA replication in S-phase, in the absence of PARP1 function, the replication fork stalls and dsDNA breaks accumulate. Given that lurbinectedin dysregulates replication fork progression, dual targeting of the replication machinery by way of PARPi has the potential to induce synergistic cell killing. The disadvantage of this approach is that PARP acts further upstream than Chk1/2 and regulates a wider range of cellular functions, increasing the risk of off-target effects. Development of PARPi has focused on their application in BRCA-mutated cancers; deficiencies in homologous repair sensitise these tumors to PARP inhibition while minimising effects on healthy cells^47–49^.

### Limitations and future experiments

While this study establishes the therapeutic potential of combining lurbinectedin, a potent and efficacious chemotherapeutic agent for treatment of SCLC, with inhibitors of key DDR kinases, questions remain regarding the precise mechanisms underpinning tumor cell responses to this combination.

Firstly, future work may need to expand on the number and diversity of SCLC cell lines versus what was utilised for this study. The four cell lines selected for the experiments detailed in this report do not encompass all transcriptional subtypes of SCLC. Given the importance of transcriptional markers in patient stratification and clinical outcomes^50,51^, assessing the replicability of the reported results across SCLC subtypes would better inform future clinical translation, and provide further insight into the relevance of the transcriptomic profile on tumor cell response to lurbinectedin and prexasertib. It would be ideal to perform further testing of this drug combination using patient-derived tissue; however, obtaining matched patient samples is complicated by the rarity of surgical intervention. One alternative is the use of circulating tumor cells derived from whole blood or pleural fluid samples to generate tumoroids, which can better recapitulate the effect of the tumor microenvironment than 2D cell culture. Tumoroids are a useful model for studying the effects of matrix mechanosensing and mechanotransduction on compound delivery^52^, a key aspect of treatment efficacy and evasion.

Dose-limiting toxicity has severely impacted the clinical potential of prexasertib, among other checkpoint kinase inhibitors. To evaluate whether our proposed combinatorial method sufficiently reduces the dose necessary to successfully target tumor cells while avoiding deleterious consequences for healthy cells, it may be useful to test our proposed treatment combination *in vivo*. Genetically engineered mouse models (GEMMs) recapitulating the *TP53* and *RB1* mutations found in SCLC provide a reasonable platform for examining the impact of these alterations on treatment outcomes^53^.

Our work has demonstrated the utility of targeting the DNA damage repair pathway to potentiate the cytotoxic effects of lurbinectedin in small cell lung cancer. We have identified key mechanisms and markers involved in treatment response. Further investigation into the temporal response and the impact of SCLC subtype may further aid in the preclinical development of this treatment modality. Future work could calculate appropriate dose regimens and patient stratification criteria by testing the combination of lurbinectedin and prexasertib in 3D and *in vivo* models. Nonetheless and despite stated limitations, the results provide a preclinical mechanistic rationale for overcoming a pro-survival drug-resistance induced G2/M cell cycle checkpoint pathway engaged by the FDA-approved lurbinectedin treatment of SCLC.

## Methods

### Cell culture

SCLC cell lines were obtained from American Type Culture Collection (ATCC) and cultured according to ATCC guidelines for each cell line. NCI-H1882, NCI-H1048 and NCI-H1105 were cultured in DMEM/F12-derived (Sigma Alrdich) HITES complete media plus 5% FBS; NCI-H526 was cultured in RPMI (Sigma Aldrich) plus 10% FBS. All media was supplemented with penicillin-streptomycin and L-glutamine.

### Chemical compounds

Prexasertib and lurbinectedin were purchased from MedChemExpress. Drugs were reconstituted in DMSO and stored at -20°C. Serial dilutions were performed in culture media corresponding to the cell type used.

### Cytotoxicity analysis by CellTiterGlo for IC50 and synergy calculations

Cells were plated at a density of 2x10^3^ and allowed to adhere for 16 hr. Cells were treated with lurbinectedin, prexasertib, or combination and incubated for 72 hr. CellTiterGlo reagent (Promega) was administered to each well. Resultant luminescent output was measured and used to evaluate cell viability relative to control wells. GraphPad Prism software was used to calculate IC50 for each drug using the cell viability data. For plates treated with lurbinectedin/prexasertib combination, the cell viability data was input to SynergyFinder. Synergy between lurbinectedin and prexasertib was calculated using the HSA reference model.

### Protein separation and qualification by gel electrophoresis and Western blot

Cells were seeded at a density of 4x10^5^ per well and allowed to adhere for 16 hr. Cells were then treated with lurbinectedin, prexasertib, combination, or DMSO. Cells were harvested and lysed after 24 hr incubation to collect protein content. Total protein content in each sample was quantified by BCA assay.

Proteins were separated by gel electrophoresis in MES running buffer (Invitrogen) at 120 V for 120 mins. Sample proteins were transferred from the gel onto ImmunoBlot membrane in tris-glycine buffer. Membrane was trimmed and blocked with 5% milk in TBST. The membrane strips were incubated in primary antibody overnight at 4°C before washing and probing with secondary antibodies. Probed membranes were imaged using the SynGene PXi system. For target proteins of the same size, membranes were stripped and re-probed following the steps above. Antibodies were obtained from Cell Signaling Technologies, Sigma, and Invitrogen.

### SubG1 analysis by propidium iodide staining and flow cytometry

H1048 cells were synchronised by culturing in serum-deprived media (HITES + 0.5% FBS) for 48 hr. The cell media was then replaced by standard HITES plus 5% FBS and treated with lurbinectedin, prexasertib, combination or DMSO control. Cells were collected after 24 hr, washed in PBS, and fixed in 70% EtOH for 24 hr. After fixation, cells were pelleted by high-speed centrifugation and washed in phosphate-citrate buffer and centrifuged again. The cell pellet was then suspended in propidium iodide plus RNase A and incubated for 30 mins at room temperature. The prepared sample was immediately analysed by flow cytometry using the CytoFLEX system (Beckman Coulter).

Statistical analysis and graphing were performed on GraphPad prism. Datapoints were expressed as mean ± SEM.

## Acknowledgements

W.S.E-D. is an American Cancer Society Research Professor and is supported by the Mencoff Family University Professorship at Brown University. This work was previously presented in apart at the annual American Association for Cancer Research meeting in March 2024 (Ashley Sanchez Sevilla Uruchurtu, Tyler Roady, Agenxnie Anderson, Wafik S. El-Deiry. Chk1/2 inhibition enhances response to lurbinectedin treatment in small cell lung cancer [abstract]. In: Proceedings of the American Association for Cancer Research Annual Meeting 2024; Part 1 (Regular Abstracts); 2024 Apr 5-10; San Diego, CA. Philadelphia (PA): AACR; Cancer Res 2024;84(6_Suppl):Abstract nr 3191.). The work by A.F.S.S.U. was conducted in partial fulfilment of the requirements for the degree of Doctor of Philosophy in the Pathobiology Graduate Program at Brown University. The authors thank thesis committee members Ian Wong, Patrycja Dubielecka, Sean Lawler, and C.G.A. at Brown University and Patrick C. Ma at Louisiana State University (LSU) Health Sciences Center for reading the manuscript and providing helpful input.

---

**Supplementary Figure 1.**
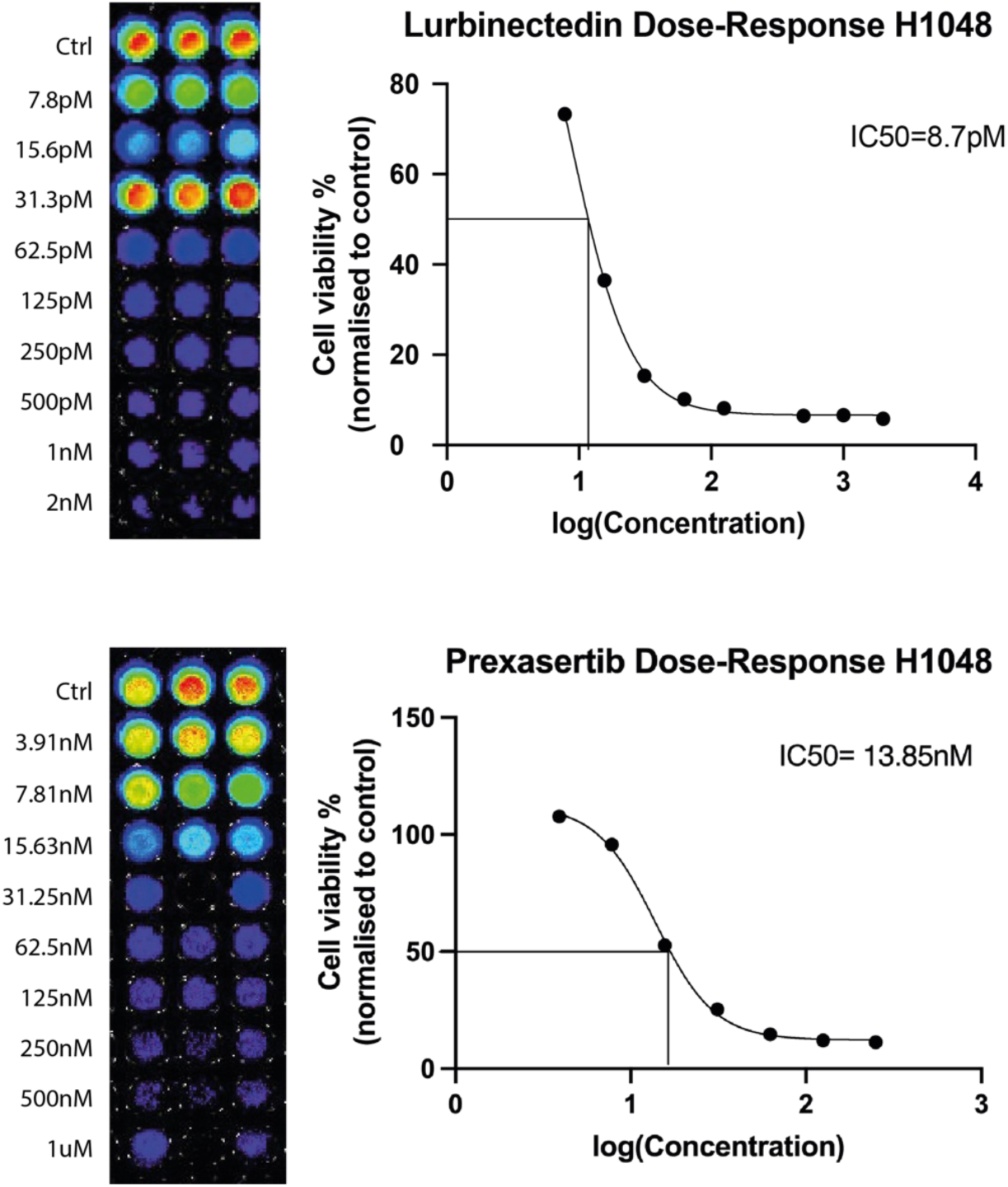
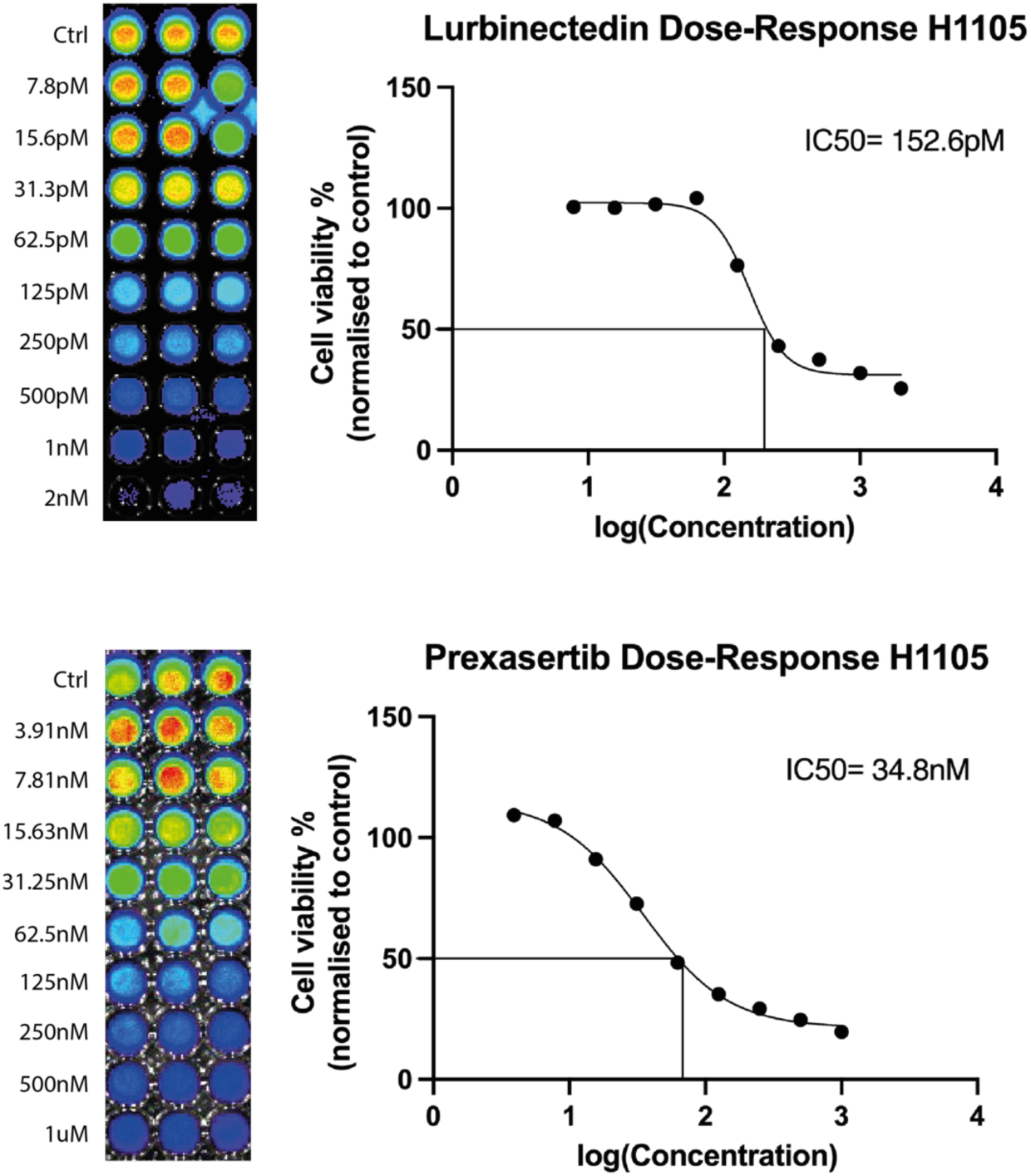
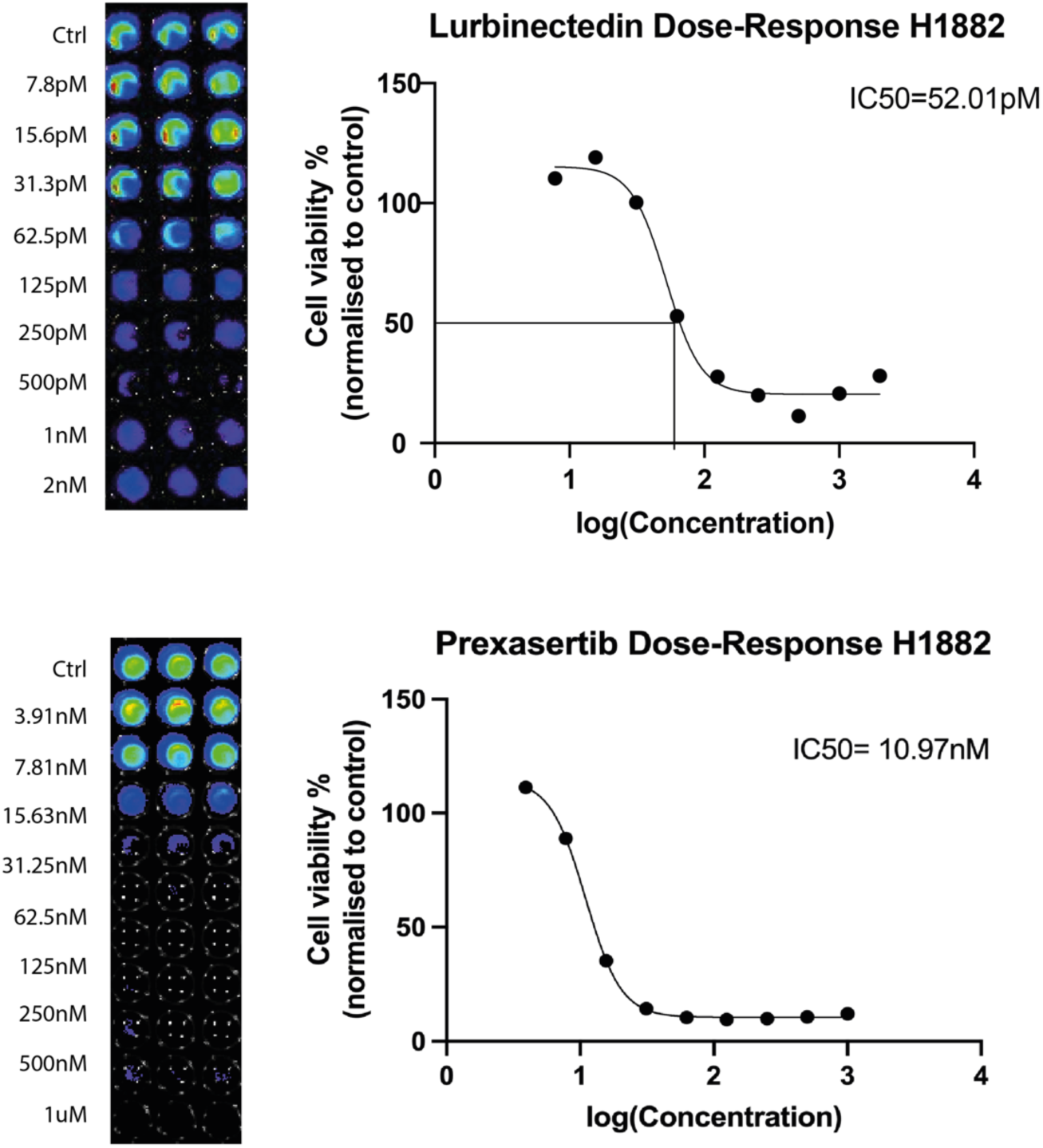
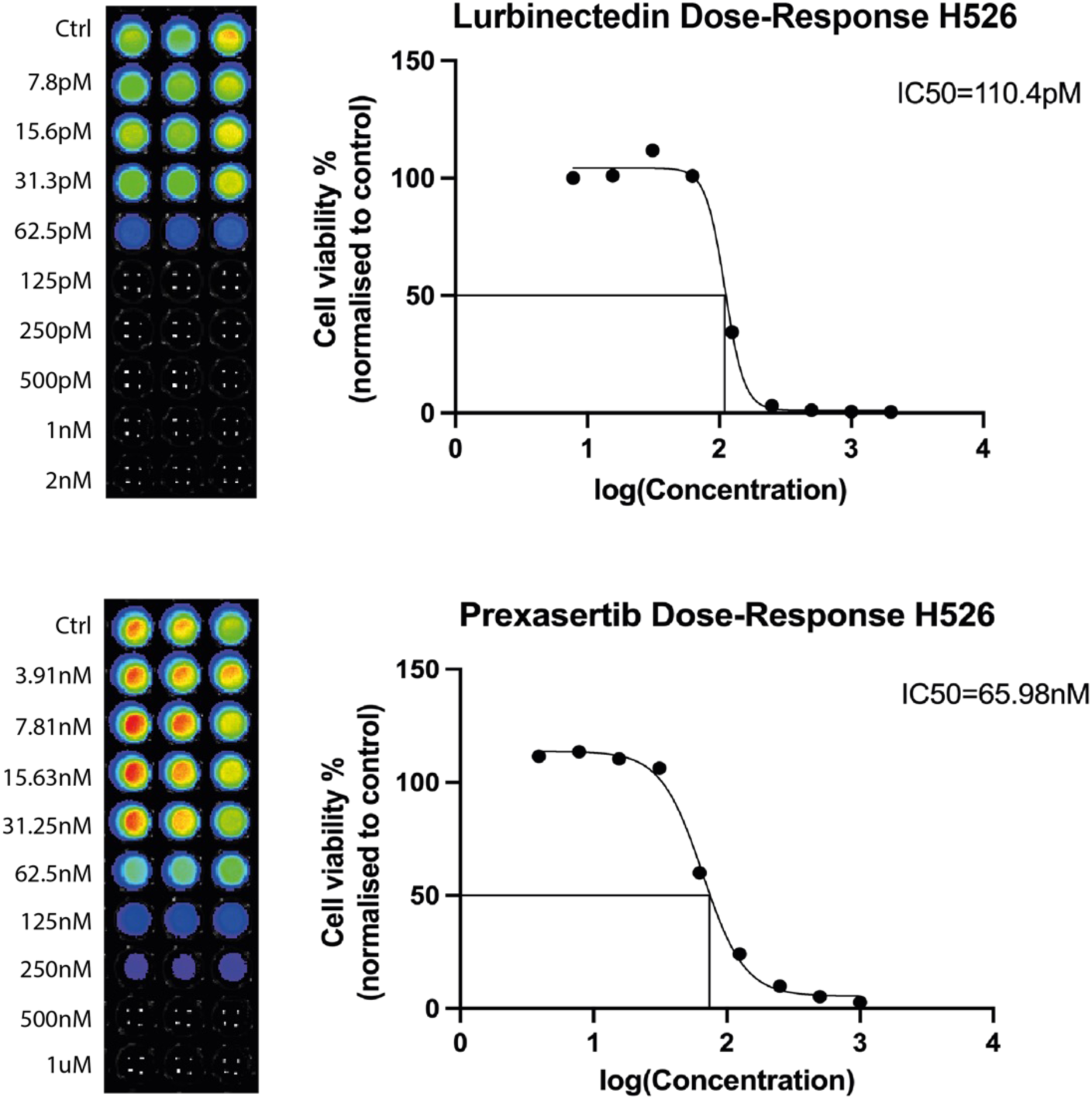
IC50 values for lurbinectedin and prexasertib were calculated for each cell line by cell viability analysis. A-D: SCLC cell lines (NCI-H1048, NCI-H1105, NCI-H1882, NCI-H526) were treated with lurbinectedin and/or prexasertib at serial doses. Cell viability was assessed at 72h by CellTiter Glo. Data was entered into GraphPad Prism software to calculate IC50 doses for each drug by cell line.

**Supplementary Figure 2.**
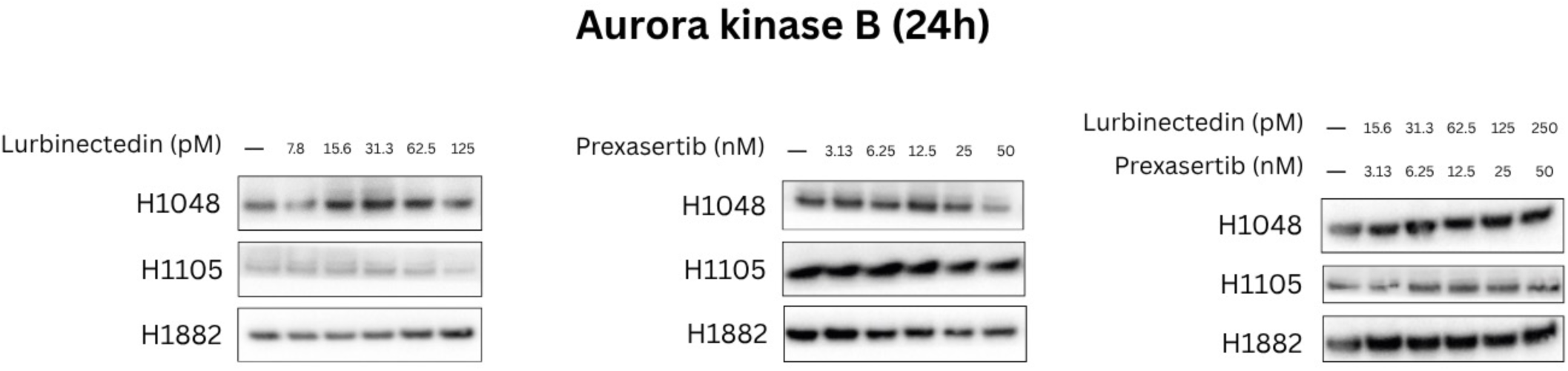
Lurbinectedin increases intracellular levels of serine-threonine kinase Aurora B in a dose-dependent manner. Conversely, prexasertib abrogates levels of Aurora. **B.** SCLC cell lines (NCI-H1048, NCI-H1105, NCI-H1882) were treated with lurbinectedin and/or prexasertib for 24h. Cells were then harvested and their protein content extracted for analysis of Aurora kinase B levels by immunoblotting.

